# pUdOs: concise plasmids for bacterial and mammalian cells

**DOI:** 10.1101/2023.07.05.547852

**Authors:** France Ourida Manigat, Louise B. Connell, Brittany N. Stewart, Abdel-Rahman LePabic, Christian J.G. Tessier, Johnathon R. Emlaw, Nicholas D. Calvert, Anthony Rössl, Adam J. Shuhendler, Corrie J.B. daCosta, François-Xavier Campbell-Valois

**Affiliations:** Center for Chemical and Synthetic Biology, Department of Chemistry and Biomolecular Sciences, University of Ottawa, Ottawa, ON K1N 6N5, Canada; bioGARAGE, Faculty of Science, University of Ottawa, Ottawa, ON, Canada; University of Ottawa Heart Institute, Ottawa, ON K1Y 4W7, Canada; Centre for Infection, Immunity and Inflammation, Department of Biochemistry, Microbiology and Immunology, University of Ottawa, Ottawa, ON K1N 6N5, Canada

**Keywords:** Plasmid, Golden Gate, proteobacteria, *Escherichia coli*, *Shigella*, mammalian cell expression

## Abstract

The pUdOs are 28 plasmids of small size combining four different origins of replication and seven selection markers, which together afford flexible use in *Escherichia coli* and several related gram-negative bacteria. The promoterless multicloning site is insulated from upstream spurious promoters by strong transcription terminators, and contains type IIP or IIS restriction sites for conventional or Golden-gate cloning. pUdOs can be converted into efficient expression vectors through the insertion of a promoter at the user’s discretion. For example, we demonstrate the utility of pUdOs as the backbone for an improved version of a Type III Secretion System reporter in *Shigella*. In addition, we derive a series of pUdO-based mammalian expression vectors affording distinct levels of expression and transfection efficiencies comparable to commonly used mammalian expression plasmids. Thus, pUdOs could advantageously replace traditional plasmids in a wide variety of cell types and applications.

## Introduction

Plasmids are circular double-stranded DNA molecules that replicate autonomously. They are frequently encountered in wild bacteria, although they are usually not essential for survival. Plasmids allow bacteria to exchange genetic material, thereby conferring evolutionary advantages, for example through genes encoding antibiotic resistance, interbacterial competition factors, or virulence factors [1, 2]. The molecular weight of natural plasmids varies from 1,000 to 400,000 base pairs and is often inflated by non-coding DNA [3].

Engineered plasmids are used for gene cloning and protein expression. They are minimally composed of an origin of replication, a selection marker, and a multiple cloning site, which are necessary for their manipulation *in vitro,* as well as their maintenance in bacteria. The origin of replication regulates the initiation of DNA replication and thus plays a critical role in maintaining the plasmid and determining its copy number per cell [4]. Plasmids harboring origins of replication from different incompatibility groups can be maintained together in the same cell [5]. The selection marker is a gene that forces cells to retain a plasmid in order to be viable, for example, in the presence of an antibiotic, or in the absence of an essential nutrient. This facilitates isolation of cells harboring the plasmid through counterselection of those that lack it. Finally, the multiple cloning site comprises several endonuclease restriction sites that enable the insertion of genetic material of interest into the plasmid. This results in a recombinant plasmid that replicates similarly to the original plasmid, allowing the replication, expression, storage, and distribution of the introduced genetic material. In eukaryotes, such as yeast, insects, plants, and mammalian cells, most engineered plasmids function as shuttle plasmids, typically manipulated in *Escherichia coli* in much the same way as described above, but with additional features required for their use in the intended eukaryotic host.

Historically, engineered plasmids have been assembled from DNA fragments obtained through restriction enzyme digestion. As a result, they typically contain excess non-coding DNA, making them larger than necessary (>3.5 kbp in most cases). Excess DNA introduces limitations, as smaller plasmids would allow the cloning of larger genetic fragments. In addition, since transformation efficiency is proportional to plasmid size, excess non-coding DNA in plasmids hinders the generation of large DNA libraries required for protein engineering, synthetic biology, and biotechnology applications [6]. Despite these limitations, few efforts have been made to minimize non-coding DNA in engineered plasmids. To this end, we introduce “plasmids from the Université d’Ottawa” (pUdO), which are a set of 28 plasmids of remarkably small size (1.2-3.2 kb), each containing only essential elements. The pUdOs were assembled by shuffling four origins of replication with seven antibiotic resistance genes. We demonstrate the utility of pUdOs by creating an improved version of the transcription-based secretion activity reporter of the Type III Secretion System in *Shigella flexneri* [7], and further by deriving plasmids for the expression of proteins in cultured mammalian cells [8].

## Results and Discussion

### pUdO Design

The pUdOs were assembled from four parts (Figure 1A), two of which were variable and corresponded to one of the four origins of replication (purple) and of the seven antibiotic resistance genes/selection markers (red). Each origin of replication and selection marker was isolated by PCR from source plasmids (Table S1 and S2). The two invariable parts were obtained through gene synthesis and constitute the multiple cloning site (blue), and the bridge (green). The multiple cloning site in each pUdO includes type IIP and type IIS restriction sites, amenable to both restriction and Golden Gate cloning. The multiple cloning site is also flanked by strong unidirectional transcriptional terminators, as well as universal primer-binding sites to facilitate sequencing (Figure S1). The bridge includes a bidirectional transcription terminator, and is flanked by two universal primer-binding sites to further facilitate sequencing (Figure S1). Each pUdO was constructed through isothermal assembly by first stitching the multiple cloning site to the antibiotic resistance gene/selection marker, which was then stitched to the bridge, and finally to the origin of replication (Figure 1B). The first set of plasmids: pUdO1a, 2a, 3a and 4a were obtained by the assembly of the four fragments described above. The rest of the set was derived from pUdO1a-4a by exchanging the resistance gene through a two-fragment assembly (Table 1). In total, the pUdOs constitute a set of 28 plasmids, ranging in size from 1193 bp to 3241 bp (Figure 1B). The small size of pUdOs results from the pruning of superfluous DNA, which is prevalent in the original source plasmids but is discarded during the amplification of individual components. Indeed, the pUdO1t and pUdO1z are among some of the smallest plasmids ever made [9].

**Figure 1.**
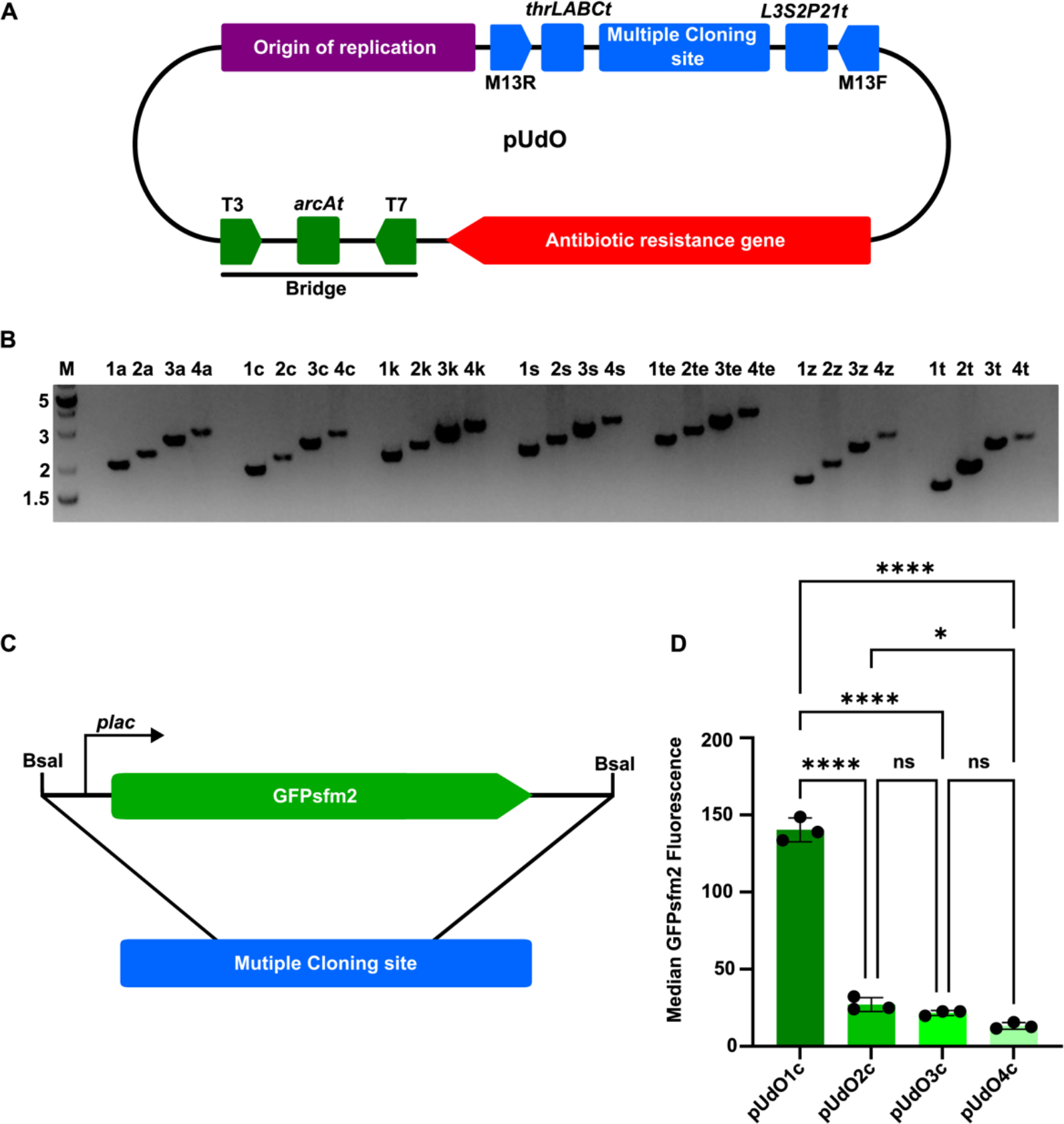
Design, construction, and validation of pUdO plasmids. (A) General map of pUdO plasmids. *thrLABCt, arcAt, L3S2P21t*: terminators; M13R, M13F, T7, T3: universal primers. (B) Validation of pUdO plasmids by restriction digestion with EcoRI. Lane M: DNA ladder; Lane 1 to 28: digested plasmids. Ladder size in kilobase pairs (kbp). (C) Insertion of *lacp*::GFPsfm2 into pUdO#c. (D) Quantification of the fluorescence produced by *lacp*::GFPsfm2 in the pUdO#c by flow cytometry, N=3. Median RFU and SD are shown here. Statistical analysis: one-way ANOVA with a 95% confidence interval and Tukey’s multiple comparison test (*p< 0.05; **p< 0.01; ***p< 0.001; ****p< 0.0001); N=3.

**Table 1.** Properties of basic and TSAR plasmids generated in this work.

| **Name** | **Origin** | **Antibiotic resistance** | **Total Size (bp)** |
| --- | --- | --- | --- |
| pUdO1a | pUC | Ampicillin | 2043 |
| pUdO2a | p15A | Ampicillin | 2262 |
| pUdO3a | pSC101 | Ampicillin | 2635 |
| pUdO4a | pBBR1 | Ampicillin | 2847 |
| pUdO1c | pUC | Chloramphenicol | 1867 |
| pUdO2c | p15A | Chloramphenicol | 2086 |
| pUdO3c | pSC101 | Chloramphenicol | 2459 |
| pUdO4c | pBBR1 | Chloramphenicol | 2671 |
| pUdO1k | pUC | Kanamycin | 2007 |
| pUdO2k | p15A | Kanamycin | 2226 |
| pUdO3k | pSC101 | Kanamycin | 2599 |
| pUdO4k | pBBR1 | Kanamycin | 2811 |
| pUdO1s | pUC | Spectinomycin | 2221 |
| pUdO2s | p15A | Spectinomycin | 2440 |
| pUdO3s | pSC101 | Spectinomycin | 2813 |
| pUdO4s | pBBR1 | Spectinomycin | 3025 |
| pUdO1te | pUC | Tetracycline | 2437 |
| pUdO2te | p15A | Tetracycline | 2656 |
| pUdO3te | pSC101 | Tetracycline | 3029 |
| pUdO4te | pBBR1 | Tetracycline | 3241 |
| pUdO1z | pUC | Zeocin | 1555 |
| pUdO2z | p15A | Zeocin | 1774 |
| pUdO3z | pSC101 | Zeocin | 2147 |
| pUdO4z | pBBR1 | Zeocin | 2359 |
| pUdO1t | pUC | Trimethoprim | 1417 |
| pUdO2t | p15A | Trimethoprim | 1636 |
| pUdO3t | pSC101 | Trimethoprim | 2009 |
| pUdO4t | pBBR1 | Trimethoprim | 2221 |
| pUdOanc-a | pUC | Ampicillin | 1844 |
| pUdOanc-t | pUC | Trimethoprim | 1193 |
| pTSAR3.1t | pUC | Ampicillin | 3949 |
| pTSAR3.3t | pUC | Ampicillin | 3895 |
| pTSAR3.4t | pUC | Ampicillin | 3914 |
| pTSAR3.5t | pUC | Ampicillin | 3923 |
| pTSAR3.6t | pUC | Ampicillin | 3902 |
| pTSAR3.7t | pUC | Ampicillin | 3881 |

The multiple cloning site includes a sequence complementary to the M13-rev universal primer, a XmaI/SmaI restriction site, the transcription terminator of *thrLABC* (*thrLABCt*), the multiple cloning site itself, the synthetic transcription terminator L3S2P21, a HindIII restriction site, and the sequence complementary to the M13-fwd universal primer (Figure S1). Successful cloning is easily confirmed through sequencing using the M13-fwd and M13-rev primers. The Xma/SmaI and HindIII restriction sites can be used to exchange the terminators and the multiple cloning site region with the genetic material of interest. The selected terminators are among the strongest and least recombination prone [10]. Of note, users must include a promoter of choice for expressing their gene of interest. Given that the terminators flanking the multiple cloning site are most efficient on the forward strand, we recommend inserting the promoter on this strand. If the transcriptional insulation of the transgene is unimportant, the insert can be introduced on the reverse strand. The multiple cloning site includes 16 type IIP restriction sites, which are cleaved by some of the most common, reliable, and affordable restriction enzymes. The selection includes those specific for endonucleases recognizing 6 or 8 bp restrictions sites and generating 5′ and 3′ sticky, or blunt, ends. Two sets of compatible sticky end restriction sites were included (i.e., i. EcoRI and MfeI; ii. and XbaI, NheI and SpeI). Flanking this region are three tandems of Type IIS restriction sites that can be used for Golden Gate cloning (i.e., BsaI, BbsI and SapI). To diminish putative interference with translation of the transgene, we avoided inserting start codons at the 5′ end and introduced stop codons in the three reading frames at both ends (Figure S1).

The bridge includes a sequence complementary to the T7 universal primer, an AscI restriction site, the bidirectional transcription terminator of *arcA* (*arcAt*), an AvrII restriction site, and a sequence complementary to the T3 universal primer (Figure S1). The bidirectional *arcAt* was also inserted in the bridge fragment to alleviate the impact of spurious promoters on the expression level of genes inserted in the multiple cloning site. The AscI and AvrII restriction sites are unique in most plasmids of the pUdO series, thereby allowing the exchange of the *arcAt* with genetic material of interest, or the cloning of genetic material upstream or downstream of *arcAt*. Finally, three stop codons are present on the sense strand both upstream and downstream of the terminator.

The four selected origins of replication belong to different incompatibility groups (Table 1): group A, pMB1/pUC (1); group B, p15a (2); group C, SC101 (3); atypical, BBR1 (4), meaning multiple different pUdOs can, in principle, be maintained in the same cell. The mutated version of the MB1 origin of replication derived from pUC and found in pUdO1x maintains 500 to 700 copies per cell [11]. The 15A origin of replication in pUdO2x maintains 10 to 12 copies per cell [12]. The SC101 origin of replication used in pUdO3x maintains approximately 5 copies per cell [11]. Lastly, the BBR1 origin of replication in pUdO4x maintains approximately 5 copies per cell [13]. The first three origins of replication allowed for the maintenance of plasmids in Enterobacteriaceae, whereas BRR1 is a broad host range origin of replication that is also functional in *Bordetella*, *Vibrio*, *Rhizobium and Pseudomonas* [14]. In short, the pUdO comes with one of four origins of replication that maintains copy numbers ranging from 5 to 500 copies per cell, which can be used to tune the expression level of the gene of interest.

Together with the origin of replication, the selection markers determine the capacity of bacteria to co-maintain a combination of plasmids. To achieve flexibility in compatibility, seven selection markers were chosen: ampicillin (a), chloramphenicol (c), kanamycin (k), spectinomycin (s), tetracycline (te), zeocin (z), and trimethoprim (t). The trimethoprim and zeocin resistance genes were selected because of their small size (i.e., 237 and 375 bp, respectively). Finally, the selection marker gene was inserted on the same strand as the multiple cloning site terminators to maximize the terminators capacity to insulate the inserted genetic material from the influence of the promoter associated with the selection marker. The naming scheme of each pUdO series follows from their origin of replication (number, ‘#’) and selection marker (letter, ‘x’). For example, pUdO1a possesses the pUC origin (‘1’) and ampicillin (‘a’) resistance gene as its selection marker (Table 1).

### Validation of pUdOs

The entire sequence of each pUdO was verified by Sanger sequencing using M13R, M13F, T3 and T7 universal primers, which anneal to the multiple cloning site and the bridge. In addition, the molecular weight of each pUdO was verified by agarose gel electrophoresis, where each of the 28 plasmids was digested with the enzyme EcoRI, which recognizes a single restriction site within the multiple cloning site. As expected, each linearized pUdO migrated according to its predicted molecular weight (Figure 1B). We also evaluated the effect of the origin of replication on gene expression. *lac*p::GFPsfm2 was cloned by Golden Gate using BsaI into the pUdO#c series, and then transformed into *E. coli* (DH10B) (Fig. 1C). Using flow cytometry, the fluorescence (mean ± standard deviation) in strains harboring the following plasmids was measured: pUdO1c (130.4 ± 7.8 RFU), pUdO2c (27.1 ± 4.5 RFU), pUdO3c (21.6 ± 1.6 RFU) and pUdO4c (13.2 ± 2.1 RFU)

(Figure 1D). This data indicated that with the pUdO#c series it is possible to modulate the level of expression 10 fold, based solely on the choice of origin of replication. Furthermore, origins of replication 2, 3, and 4 all afforded similar levels of expression, which could prove useful when seeking to express components at near equivalent molar or stoichiometric ratios.

### Improved Type III Secretion Reporter based on the pUdOs

Strong terminators were included in pUdOs to facilitate the construction of synthetic systems containing multiple cistrons, and with minimal transcriptional interference from spurious promoters. To assess this aspect of the pUdOs, we chose the transcription-based secretion activity reporter (TSAR) system as a case study. The TSAR system consists of an *ipaH7.8*p::GFPsfm2 (hereafter GFP) reporter gene, which is upregulated when the Type Three Secretion System (T3SS) of *Shigella* is active [7]. This upregulation depends on the transcription activator MxiE [15]. In addition, the TSAR system is combined with a constitutively expressed *rpsM*p::fluorescent protein reporter gene (Figure 2A). We derived a new generation of pTSAR plasmids through the insertion of the two corresponding reporter genes from pTSAR1ud2.4s [7] and the L3S3P21 terminator into the pUdO1a backbone, yielding pTSAR3.4t. From this plasmid, we derived pTSAR3.1t (mCerulean), 3.3t (DsRed T3 S4T), 3.5t (eBFP2), 3.6t (mScarlet), and 3.7t (E2- Crimson). These fluorescent proteins were selected because they covered a wide range of excitation and emission wavelengths [16–20], which are compatible when used in combination with GFP. To confirm that the pTSAR3.#t retained the original T3SS-regulated temporal expression, HeLa cells were coinfected with *Shigella flexneri* wild-type and Δ*mxiE* strains carrying different pTSAR3xt derivatives, which allowed strain discrimination by fluorescence microscopy. Consistent with previous findings [7], wild-type but not Δ*mxiE* displayed GFP fluorescence 60- minute post-infection when located inside epithelial cells (Figure 2B-2F), whereas no GFP signal was observed in bacteria located outside of cells. These observations indicated that the TSAR signal is robust following invasion of cultured cells with *Shigella* harboring pTSAR3.#t as reported with the first generation of pTSAR [7].

**Figure 2.**
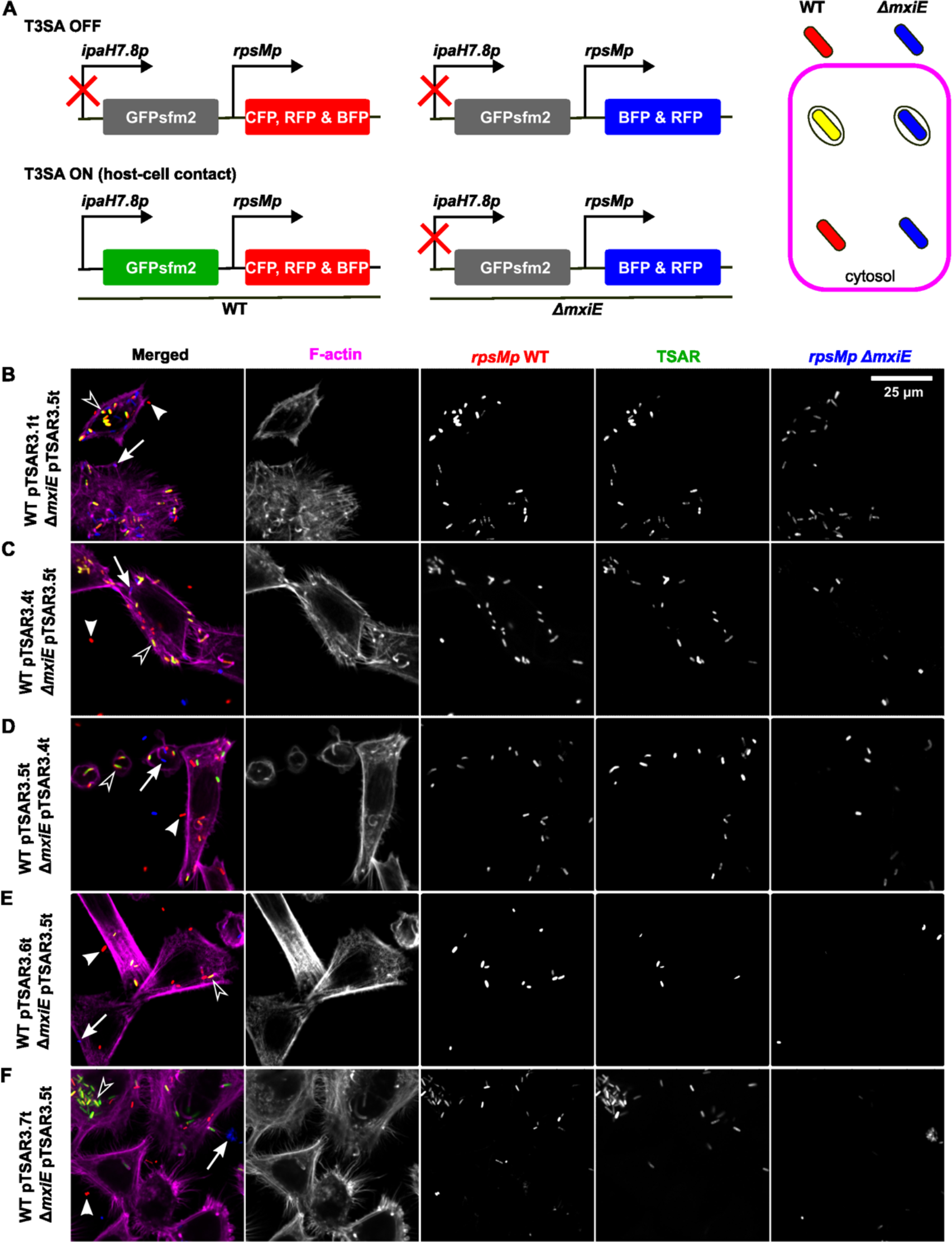
pTSAR3.#t discriminates between intracellular and extracellular bacteria. (A) Schematic of pTSAR reporter. Micrographs of Hela cells monolayer co-infected with (B) *Shigella flexneri* strains wild-type harboring pTSAR3.1t (mCerulean) and *ΔmxiE* pTSAR3.4t (mcherry), (C) wild-type pTSAR3.4t (mCherry) and *ΔmxiE* pTSAR3.5t (eBFP2), (D) wild-type pTSAR3.5t (eBFP2) and *ΔmxiE* pTSAR3.4t (mCherry), (E) wild-type pTSAR3.6t (mScarlet) and *ΔmxiE* pTSAR3.5t (eBFP2) and, (F) wild-type pTSAR3.7t (e2Crimson) and *ΔmxiE* pTSAR3.5t (eBFP2) respectively and counterstained with Phalloidin-Alexa Fluor 647. Arrows point to actively secreting intracellular bacteria, arrowheads and hollow arrowheads point respectively to intracellular and extracellular non-secreting bacteria.

Due to the excitation wavelength of the DsRed, pTSAR1.3 and its equivalent pTSAR3.3t are well suited for most flow cytometers (Figure 3A). To compare the performance of pTSAR1.3 and its equivalent pTSAR3.3t, we used flow cytometry to measure the respective median fluorescence intensities of the TSAR (GFP) and the *rpsM*p::DsRed in Δ*mxiE* and in Δ*ipaD*, which displays minimum and maximum TSAR signals *in vitro* due to the loss of the transcription activator MxiE or the constitutive activity of the T3SS, respectively [7] . As expected, minimal TSAR signal was observed in Δ*mxiE*, whereas in Δ*ipaD*, a strong TSAR signal was observed in cells harboring pTSAR1.3 (49.5 ± 4.9 RFU) or pTSAR3.3t (188.3 ± 5.7 RFU), yielding respectively 165- and 471-fold increase in the signal-to-noise ratio (Δ*ipaD*/Δ*mxiE*) (Figure 3B, Figure S2). Previously, we observed an artifactual increase in expression of the DsRed located downstream of the TSAR when the latter was active [7]. Indeed, the DsRed produced by pTSAR1.3 in Δ*ipaD* (19.1 ± 0.5 RFU) nearly doubled that in Δ*mxiE* (10.7 ± 0.9), whereas the same strains harboring pTSAR3.3t yielded similar fluorescence (17.9 ± 1.7 to 19.5 ± 2.3, respectively) (Figure 3B, S2). In short, the pTSAR3.3t provides a stronger TSAR signal-to-noise ratio while at the same time being less prone to interfere with the expression of the constitutively expressed DsRed than its pTSAR1.3 predecessor. The inclusion of stronger terminators in pTSAR3.3t likely explains the increased insulation of the DsRed cistron from the influence of the TSAR. The increased GFP signal of pTSAR3.3t could stem from at least two factors. First, the Shine Dalgarno of the GFP used in pTSAR3.3t is more efficient [7]. Second, the addition of a stronger terminator may yield a shorter 3’ UTR that would stabilize the mRNA by preventing a decay mechanism triggered by stalled ribosomes [21]. Finally, it is noteworthy that these plasmids differ in the respective orientation of the two cistrons (Figure 3A). This as well as other variations in the assembly of the two constructs could also explain the improved performance of pTSAR3.3t.

**Figure 3.**
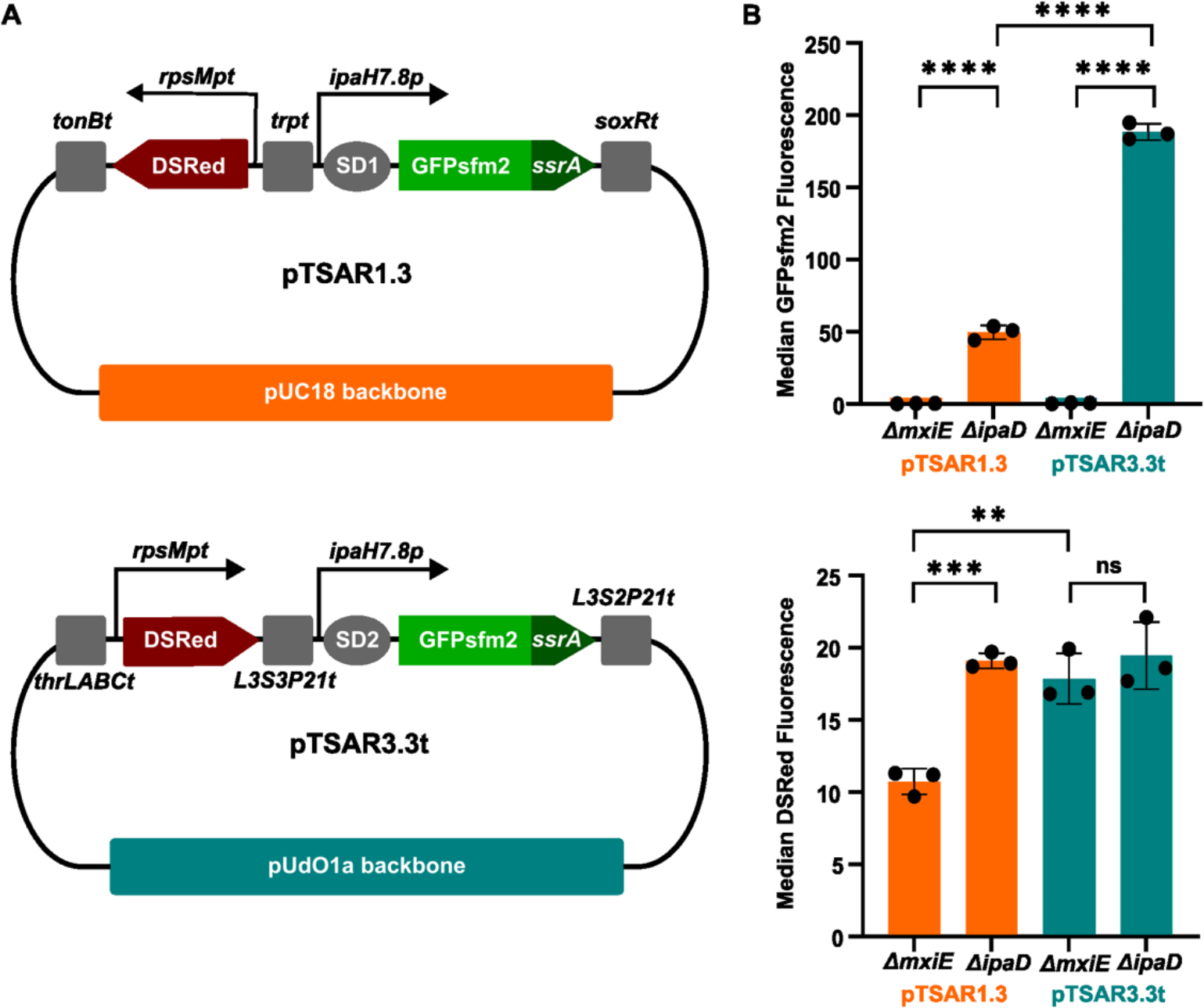
pTSAR3.3t measures Type III Secretion System activity with increased sensitivity without impacting the constitutive expression of the DsRed. (A) Maps of pTSAR1.3 and pTSAR3.3t. (B) Measurement of GFPsfm2 and DsRed fluorescent signal of pTSAR1.3 and pTSAR3.3t by flow cytometry. Median RFU, SD, and one-way ANOVA with a 95% confidence interval and Tukey’s multiple comparison test are shown here. (*p< 0.05; **p< 0.01; ***p< 0.001; ****p< 0.0001); N=3.

We next looked at the performance of pTSAR3.4t in confocal microscopy, using its predecessor pTSAR1ud2.4s as a benchmark (Figure 4A). We infected HeLa cells with wild-type *S. flexneri* carrying either pTSAR1ud2.4s or pTSAR3.4t (Figure 4B). To compare their performance, we measured pixel intensities of the fluorescent signal for both the mCherry and the GFP for approximately 200 bacteria in triplicate. We observed a higher GFP signal (1.6-fold increase), in line with that obtained in flow cytometry, suggesting the new generation pTSAR3.4t can detect the activity of the T3SS with increased sensitivity (Figure 4C). As the latter and its predecessor pTSAR1ud2.4s used the same Shine-Dalgarno for the GFP, it suggests that the terminators are the key element that makes pTSAR3.4t brighter. In contrast with the flow cytometry data, there was no significant difference in the mCherry signals of pTSAR3.4.t and pTSAR1ud2.4s. This discrepancy might stem from the differences in the construction of pTSAR 1.3 and pTSAR1ud2.4s (Figure 3A and 4A) [7], from the experimental setup, or both. In the flow cytometry experiments, we studied *Shigella* populations that were more homogenous than in the microscopy experiments. Indeed, the τι*ipaD* is a constitutively secreting strain that displays a homogeneously high TSAR signal, whereas the τι*mxiE* lacking the transcription activator required for activating the expression of the GFP display a homogeneously low TSAR signal. By contrast, the WT strain used in the microscopy experiments forms a heterogenous population as bacteria are uncoordinated during the infection of the human cell monolayer. Thus, individual bacteria may show different levels of fluorescence depending on the time elapsed since they last activated their T3SS [7]. Nevertheless, the pTSAR3.#t series is better constructed, and performs better than the original system. In the future, the improved pTSAR3.#t series should be used to monitor the activity of the T3SS in *Shigella*.

**Figure 4.**
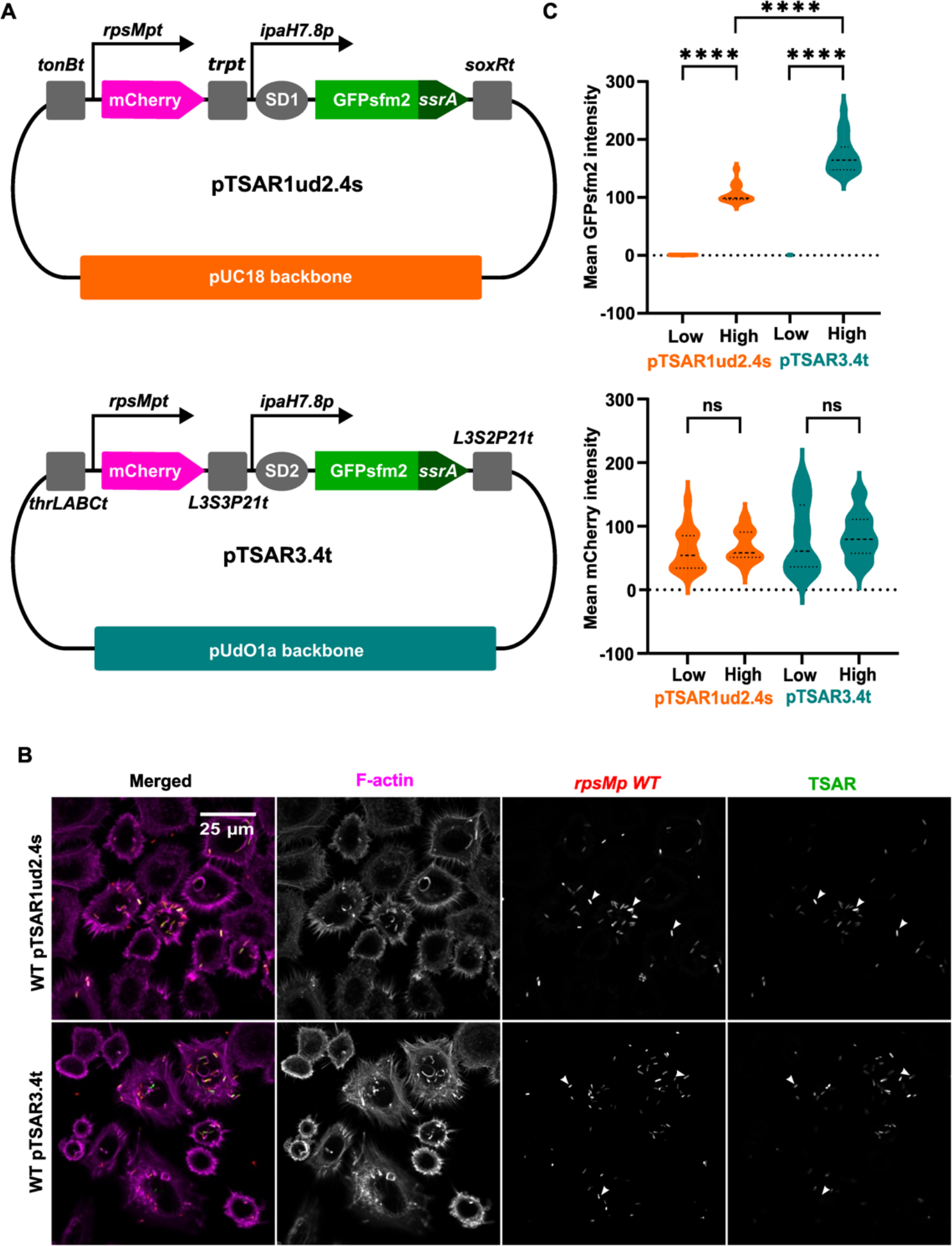
pTSAR3.4t measures Type III Secretion System activity with increased sensitivity by fluorescence microscopy. (A) maps of pTSAR1ud2.4s and pTSAR3.4t. (B) Micrographs of HeLa cell monolayer infected with wild type (WT) *Shigella flexneri* harboring pTSAR1ud2.4s or pTSAR3.4t. Arrows point to bacteria with high GFP expression. (C) Quantification of GFP and RFP fluorescence. Mean pixel intensities were measured with FIJI and statistical significance was tested with one-way ANOVA and Tukey’s multiple comparison test (*p< 0.05; **p< 0.01; ***p< 0.001; ****p< 0.0001). N=3.

### pUdOm: pUdOs for expression in mammalian cells

To expand the utility of pUdOs, we made a set of sixteen mammalian expression vectors (pUdOm#.#) derived from the original pUdO backbones by flanking the multiple cloning site with a mammalian-specific transcriptional promoter and terminator (Figure 5; Table 2). For the promoter, we chose the well-characterized cytomegalovirus (CMV) promoter and enhancer sequence, which yields a robust expression in several mammalian cell lines [22–24]. The cytomegalovirus promoter/enhancer sequence was inserted upstream of the original pUdO multiple cloning site (Figure 5A). To expand the dynamic range of expression possible with pUdOms, we made several additional plasmids with truncations to the CMV promoter/enhancer sequence (Figure 5A; Table 2) [25–28]. Finally, to terminate the transcription, we inserted either a bovine growth hormone (BGH; pUdOm#odd.#) or a simian virus 40 (SV40; pUdOm#even.#) polyadenylation signal downstream of the multiple cloning site (Figure 5A). These two established transcriptional terminators are used in a variety of mammalian expression vectors [29, 30], with the BGH terminator used in the popular pcDNA3.1. Note that the original pUdOms do not contain a 5’UTR, so the user can insert a 5’UTR of their choice during cloning. For convenience, we also provide pUdOm1.0 and pUdOm2.0 that carry a 5’UTR and an SP6 universal primer upstream of the multiple cloning site (Table 2).

**Figure 5.**
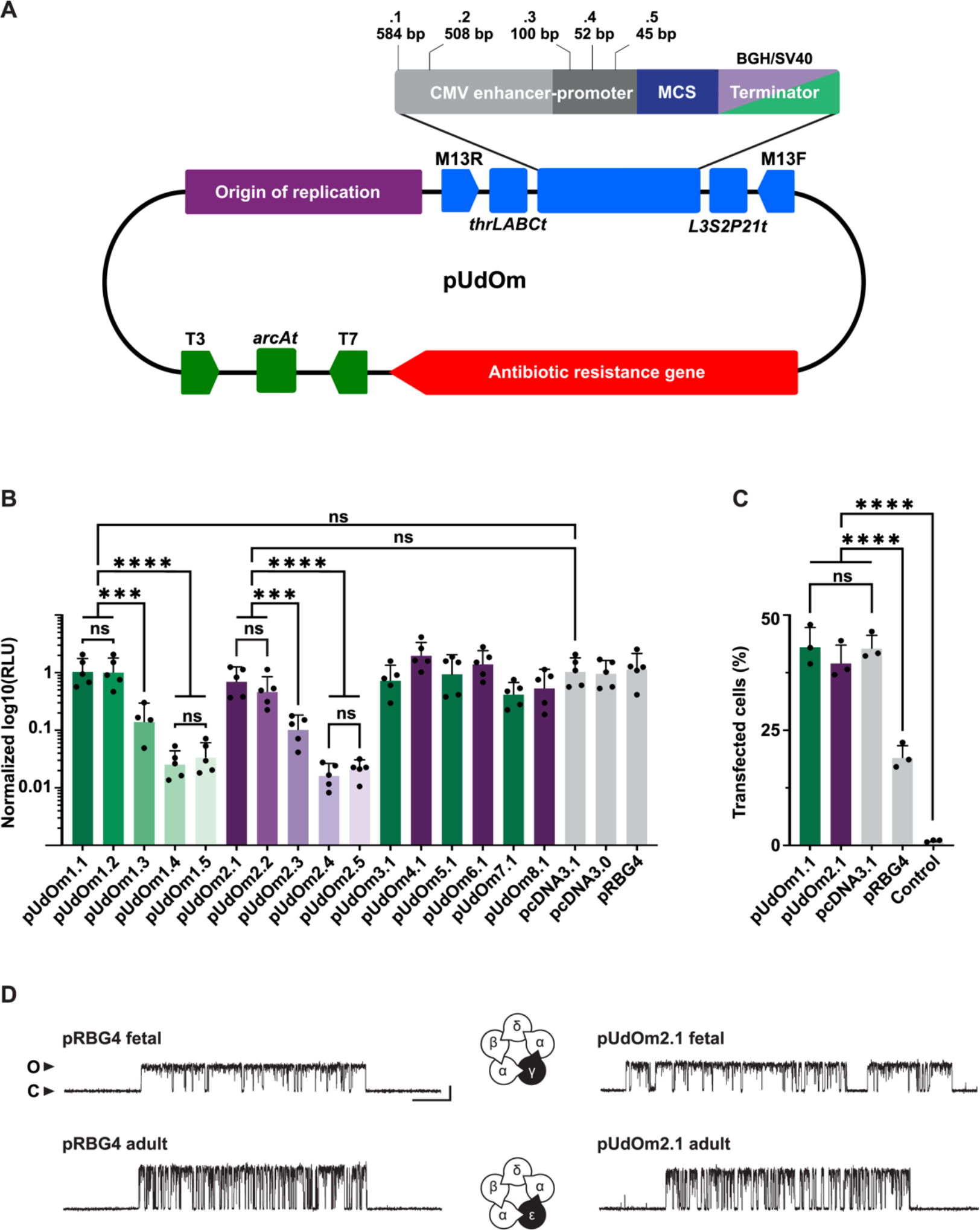
Mammalian pUdO plasmids are comparable to traditional mammalian expression vectors. (A) General map of pUdOms derived from pUdO1a (ampicillin), pUdO1c (chloramphenicol), pUdO1t (trimethoprim), and pUdO1z (zeocin) (Table 2). The ampicillin-resistant pUdOm was further expanded by truncating the CMV promoter (dark grey) and enhancer (light grey) regions (length of CMV promoter and enhancer is shown for the truncated constructs). Finally, plasmids contained a multiple cloning site (MCS; blue) and a BGH (green) or SV40 (purple) terminator. (B) HEK293 cells were transfected with the indicated pUdOm containing firefly luciferase. Luminescence was measured 24 hrs post transfection (N=4 or 5). Luminescence (RLU) was normalized to that of pcDNA3.1. Significance values were calculated by performing a one-way ANOVA and Tukey’s multiple comparison test (*p< 0.05; **p< 0.01; ***p< 0.001; ****p< 0.0001) and are presented in *Supplementary* Figure 3. (C) High-expressing pUdO plasmids have the same transfection efficiency as pcDNA3.1. Percent transfection was calculated by dividing the number of GFP-positive cells by the total number of counted cells. Bars are mean ± one standard deviation (error bars) for (points; N=3 transfections, with up to 10,000 cells counted per sample). Statistical significance was tested by performing a one-way ANOVA and Tukey’s multiple comparison test (*p< 0.05; **p< 0.01; ***p< 0.001; ****p< 0.0001); N = 3. (D) Single-channel patch clamp recordings show that fetal (γ-subunit; top) and adult (ε-subunit; bottom) acetylcholine receptors expressed from pUdOm2.1 plasmids (right) are indistinguishable from channels expressed from pRBG4 (left). Scale bar is 25 ms and 5 pA.

**Table 2.** Properties of pUdOm derivatives for expression in mammalian cells.

| **Name** | **Derived from** | **Promoter/enhancer size (bp)** | **Terminator** | **Antibiotic resistance** | **Size (bp)** |
| --- | --- | --- | --- | --- | --- |
| pUdOm1.0 <sup>a</sup> | pUdO1a | 584 | BGH | Ampicillin | 2973 |
| pUdOm1.1 | pUdO1a | 584 | BGH | Ampicillin | 2852 |
| pUdOm1.2 | pUdO1a | 508 | BGH | Ampicillin | 2776 |
| pUdOm1.3 | pUdO1a | 100 | BGH | Ampicillin | 2368 |
| pUdOm1.4 | pUdO1a | 52 | BGH | Ampicillin | 2320 |
| pUdOm1.5 | pUdO1a | 45 | BGH | Ampicillin | 2313 |
| pUdOm2.0 <sup>a</sup> | pUdO1a | 584 | SV40 | Ampicillin | 2883 |
| pUdOm2.1 | pUdO1a | 584 | SV40 | Ampicillin | 2762 |
| pUdOm2.2 | pUdO1a | 508 | SV40 | Ampicillin | 2686 |
| pUdOm2.3 | pUdO1a | 100 | SV40 | Ampicillin | 2278 |
| pUdOm2.4 | pUdO1a | 52 | SV40 | Ampicillin | 2230 |
| pUdOm2.5 | pUdO1a | 45 | SV40 | Ampicillin | 2223 |
| pUdOm3.1 | pUdO1c | 584 | BGH | Chloramphenicol | 2676 |
| pUdOm4.1 | pUdO1c | 584 | SV40 | Chloramphenicol | 2586 |
| pUdOm5.1 | pUdO1t | 584 | BGH | Trimethoprim | 2226 |
| pUdOm6.1 | pUdO1t | 584 | SV40 | Trimethoprim | 2136 |
| pUdOm7.1 | pUdO1z | 584 | BGH | Zeocin | 2364 |
| pUdOm8.1 | pUdO1z | 584 | SV40 | Zeocin | 2274 |

<sup>a</sup> These plasmids have a 5'UTR that includes a SP6 universal primer for sequencing.

To quantify the ability of pUdOms to express protein in mammalian cells, we cloned firefly luciferase into each pUdOm before transiently transfecting them into HEK293 cells and measuring luciferase activity (Figure 5B). There was no significant difference in luciferase activity, and thus expression, between the two longest (pUdOm#.1 vs. pUdOm#.2), and between the two shortest (pUdOm#.4 vs. pUdOm#.5) CMV promoters (Figure 5B; Supplementary Figure S3). For this reason, we refer to pUdOm#.1 and pUdOm#.2 as the “high expressing” tier, and pUdOm#.4 and pUdOm#.5 as the “low expressing” tier, with pUdOm#.3 constituting the “intermediate expressing” tier. There was an approximately five to seven-fold change between each expression tier, with high-expressing plasmids (pUdOm#.1 or pUdOm#.2) giving expression levels indistinguishable from pcDNA3.1 and pcDNA3.0. Comparisons across terminators (pUdOm1.# vs. pUdOm2.#), but within the same expression tier, showed that the terminator did not affect the expression level (Supplementary Figure S3). Finally, when a different pUdO (i.e., pUdO1c, pUdO1t, and pUdO1z) was chosen as the pUdOm backbone (i.e., pUdOm3.1, 4.1, 5.1, 6.1, 7.1, and 8.1), there was little change in the overall expression level, indicating that the antibiotic resistance gene had marginal effect on expression.

To compare transfection efficiency, we used flow cytometry to determine the percentage of cells expressing GFP from pUdOm1.1 and the control pcDNA3.1 (Figure 5C). We found that the transfection efficiency of pUdOm1.1 is indistinguishable from that of the commonly used pcDNA3.1. Furthermore, replacing the BGH terminator with the SV40 terminator (pUdOm1.1 vs. pUdOm2.1) did not affect transfection efficiency (Figure 5C). To ensure that pUdOms can effectively replace traditional expression vectors, we cloned the α-, β-, δ-, ε-, and γ-subunits of the human muscle-type acetylcholine receptor into the pUdO1m2.1 backbone for single-channel patch clamp experiments. The pUdOm2.1 backbone contains the same CMV promoter and SV40 terminator as pRBG4, a vector extensively used for patch clamp studies of the human muscle-type acetylcholine receptor [31]. We found that the single-channel behavior of both adult (ε-containing) and fetal (γ-containing) human acetylcholine receptors expressed from the pUdOm2.1 backbone were indistinguishable from those expressed from pRBG4, indicating that pUdOms can effectively replace standard mammalian expression vectors.

## Conclusion

We have reported the design and application of a set of concise plasmids and demonstrated their utility in *E. coli*, *S. flexneri*, and in a mammalian cell line. Their miniature size, the use of strong terminators, and the ability to use traditional or Golden Gate cloning to insert the genetic material of interest will make pUdOs advantageous for numerous applications in molecular, synthetic, and cell biology. Flexibility is embedded in the design of the pUdOs, which allows for the facile generation of derivatives for use in other model organisms or chassis.

## METHODS

### Bacterial strains and Growth conditions

*E. coli* strains DH10B and DH5α were used as the host for plasmid propagation. The *Shigella flexneri* 5a strain M90T-Sm (GenBank CM001474) harboring the virulence plasmid pWR100 (GenBank: AL391753.1) and a streptomycin resistance mutation [32] was used as the wild-type strain. The derivative mutant strains *ΔipaD*, and *ΔmxiE* were previously described [33, 34]. *E. coli* was grown in Luria-broth (LB) (Fisher Bioreagents, BP1426-2) or 2x Yeast Extract Tryptone (2xYT). *S. flexneri* was grown in Tryptic soybean-casein digest medium, Bacto (Difco Laboratories, 211822). Solid culture plates were prepared using Luria Agar base, Miller (Difco Laboratories, 211829) and Tryptic soybean-casein digest agar, Difco (Difco Laboratories, 236920) with 0.01% Congo red. Antibiotics were used for selection where applicable at the following concentrations: ampicillin 100µg/mL, kanamycin 50 µg/mL, chloramphenicol 30 µg/mL, tetracycline 12.5µg/mL, zeocin 12.5 µg/mL, trimethoprim 50 µg/mL and spectinomycin 100µg/mL.

### Design, construction and validation of the pUdOs

Primers were generated using the NEBuilder software (New England Biolabs) using the following parameters: Gibson Assembly Master Mix, 4 or more fragments, total construct length less than 10 kb, minimum overlap length: 20 nucleotides, PCR product: Q5 HiFi DNA polymerase, PCR primer concentration: 500 nM and minimum primer length: 18 nucleotides. The suggested primers were obtained from Integrated DNA Technologies (IDT) (Table S3). The pUdO#a was obtained by assembling four fragments (Table 1 and Figure S1) using the Gibson Assembly^®^ Master Mix (New England Biolabs, E2611) according to the supplier’s protocol. The origin of replication and antibiotic resistance gene fragments were obtained from source plasmids by PCR amplification with Q5 HiFi DNA polymerase (NEB). The multiple cloning site, and the bridge, were obtained by gene synthesis (gBlocks, IDT), and similarly amplified by PCR. Our initial assemblies failed due to the presence of primer dimers in the PCR of the multiple cloning site. These resulted in plasmid lacking most of the multiple cloning site but the region corresponding to the primer binding sites. We nevertheless conserved these plasmids called pUdOanc-a and pUdOanc-t (Table 1), as they could be advantageous as cloning vectors given their small size. Yet, to obtain the sought-after full-length plasmids, we integrated changes to the Gibson assembly protocol. First, we systematically treated the PCR product corresponding to the four fragments with DpnI. All PCR products were column purified. Nevertheless, if upon agarose gel analysis, a given PCR product displayed secondary product or primer dimers, we gel-purified the desired product. To obtain a reproducible high number of colonies containing the recombinant plasmid, we electroporated a small fraction of the assembly reaction into inhouse DH10B cells using 1 mm cuvettes and a micropulser (Bio-Rad) set at 1.8 kV. The sequence of each plasmid was validated by Sanger sequencing (Genome Québec) with the sequencing primers described above. Returned sequences were then aligned using CAP3 software, and manual corrections were made using the chromatograms if necessary, or by comparing overlapping reads. The pUdO#c,k,s,te,z or t (Table 1) were derived from pUdO#a by replacing the ampicillin resistance gene with the corresponding resistance gene by two-fragment assembly using the NEBuilder HiFi DNA Assembly Master Mix (NEB E2621) according to the supplier’s protocol with the above modifications.

### Construction of the pTSAR3.#t

The pTSAR3.#t set was obtained by subcloning through isothermal assembly the *rpsMp*-mCherry- *ipaH7.8p*-GFPsfm2 fragment from pTSAR1ud2.4s [7] obtained by PCR with primers GFPsfm2_fwd and GFPsfm2_rev (Table S1). This removed most of the multiple cloning site and introduced NheI and XbaI at the 5’ and 3’ of the insertion point, respectively. The cloned insert is thus flanked by the terminators in the resulting construct. It then underwent two rounds of mutagenesis PCR to remove a sequence with partial overlap with the M13 forward universal primer located downstream of *rpsMp*. Next, we replaced the tandem Trp terminator by the strong terminator L3S3P21 [10], yielding pTSAR3.4t. Two new restriction sites, EcoRV and NheI, were introduced on either side of the new terminator. The other members of the pTSAR3.#t series were derived from pTSAR3.4t by replacing mCherry with either one of these fluorescent proteins: mCerulean (pTSAR3.1t), DsRed.T3 S4T (pTSAR3.3t), EBFP2 (pTSAR3.5t), mScarlet (pTSAR3.6t), and E2-Crimson (pTSAR3.7t) [16–20].

### Construction of the pUdOm and their derivatives

To construct the empty mammalian pUdO vector (pUdOm), an SV40 terminator was first inserted into the pUdO backbone through isothermal assembly. Then, the appropriate CMV promoter was inserted at XmaI and EcoRI sites, giving rise to pUdOm2.1 and pUdOm2.2. CMV truncations were made via PCR amplification of the entire plasmid except the portion of the CMV promoter being removed, making pUdOm2.3-5. To make the pUdOm1 series, the BGH terminator replaced the SV40 terminator in the pUdOm2.1 backbone through isothermal assembly, and then subsequently to the remainder of the SV40 series through restriction enzyme cloning using KpnI and XbaI sites. The pUdOm#.1-5 do not have a 5’UTR, and this feature should be inserted upstream of the gene of interest. However, we generated pUdOm1.0 and pUdOm2.0 by ligating at a SacI restriction site a 5’UTR derived from that of Luciferase-pcDNA3, which also contains a SP6 universal primer. Luciferase-pcDNA3 was a gift from William Kaelin (Addgene plasmid # 18964 ; http://n2t.net/addgene:18964; RRID:Addgene_18964). The firefly luciferase from this plasmid was ligated into the pUdOm at SacI and XbaI sites. SacI cuts at the end of the CMV promoter, and thus the 5’UTR was brought into the mammalian pUdO backbone with the luciferase coding sequence. GFP was cloned into pUdOm2.1 between SacI and HindIII. It was then cloned from pUdOm2.1-GFP into pcDNA3.1(+) and pUdOm1.1 at SacI and XbaI sites.

### Plasmids distribution

Addgene distributes most of the plasmids reported in this study.

### *Shigella* flow cytometry

*Shigella* strains Δ*ipaD* and Δ*mxiE* harboring pTSAR1.3 [7] and the new construct pTSAR3.3t were grown overnight at 30°C with appropriate antibiotic and then sub-cultured at a ratio of 1/100 for 3 hours at 37°C in TCS broth. Sub-cultured cells were pelleted (6000xg, 2 min, RT), washed twice with PBS and incubated in propidium iodide (PI) stain (Molecular probes, P3566) at 3 µM in 1xPBS for 10 minutes at room temperature. 20 µL of cells suspension was then diluted in 980 µL PBS and assayed. The fluorescence intensities of the samples were measured with a Gallios Flow cytometer (Beckman Coulter): BFP, Ex: 405nm Em:450/50nm; GFP, Ex:488nm Em:525/50nm; dsRED, Ex:562nm Em:582/15nm; PI, Ex:633nm Em:670/30nm) and analyzed with the software KALUZA (v.2.1). Graphs and statistical analysis were prepared with GraphPad Prism (v 9.0).

### Tissue culture and infection of HeLa cells with *Shigella*

HeLa cells were cultured in EMEM (Wisent, 320-026-CL) supplemented with 10% FBS (Multicell, 081105) and 1% Pen Strep (Wisent, 450-200-EL). Coverslips were incubated with 10 µg/mL polylysine in 1xPBS for 1h and washed three times with PBS and seeded with HeLa cells at a concentration of 2.2 x 10^5^ per well in 24-well plate. Cells were grown for 24 hours at 37°C, 5% CO2 to 80-90% confluence. Wild-type Shigella harboring pTSAR1ud2.4s, pTSAR3.1t, pTSAR3.4t, pTSAR3.5t, pTSAR3.6t and pTSAR3.7t; and *ΔmxiE* mutant harboring pTSAR1ud2.4s, pTSAR3.4t, pTSAR3.5t were grown overnight at 30°C and sub-cultured for 3 hours in TCS broth at 37°C, then collected by centrifugation (6000xg, 2 min, RT) and washed with PBS. Bacterial cells were incubated with 10 µg/mL polylysine/PBS for 15min, washed three times with 1xPBS and resuspended in EMEM, 20mM HEPES. Hela cells were washed three times with EMEM, 20 mM HEPES and infected with *Shigella* strains of choice at a MOI of 50. Infected cells were allowed to incubate at room temperature for 10 min, the infecting medium was then removed and replaced with EMEM, 20 mM HEPES, then incubated at 37° with 5% CO2 for 30 min. Coverslips were then washed three times with PBS and EMEM (10% FBS, 20 mM HEPES), 50 μg/mL gentamicin was added, and the cells were incubated at 37° with 5% CO2 for an additional 30 min.

### Imaging of *Shigella* infection

Coverslips containing infected HeLa cells were washed twice with 1xPBS and fixed with 4% PFA. Cells were permeabilized in 1xPBS, 0.1% Triton X100 and blocked in 1 x PBS, 1% BSA (30min, RT). Coverslips were washed three times with 1 x PBS and incubated with phalloidin diluted in

1xPBS, (20min, RT), and then DAPI diluted in 1xPBS, 1% BSA (15 min, RT). Stained coverslips were then washed three times with 1x PBS and mounted on microscope slides (VWR) with homemade MOWIOL mounting medium. Samples were imaged with a 63X/1.46 NA oil objective on a Zeiss LSM880 confocal microscope. Images were analyzed with FIJI [35]. The mean pixel intensity of bacteria in the GFP (TSAR) and mCherry (*rpsM*p) channels were measured. The channels were split, and the RFP channel was used to auto-threshold the image using the MaxEntropy setting and then converted into a binary mask that was next used to quantify the pixel intensity of the original image. Objects with areas smaller than 0.5 μm^2^ or larger than 7 μm^2^ were discarded.

### Mammalian gene expression assays

HEK293 cells were grown in 24-well plates in Dulbecco’s modified Eagle medium supplemented with 10% fetal bovine serum and 1% penicillin/streptomycin in a 5% CO2 incubator at 37 °C. Cells were transfected with 500 ng DNA at approximately 70% confluence using ViaFect transfection reagent (Promega) and assayed 24 hours after transfection. For luciferase assays, cells were lysed using 5X Passive Lysis Buffer (Promega) and assayed using the ONE-Glo EX luciferase assay system (Promega) on a SpectraMax L luminometer (Molecular Devices) with a 0.5 s integration time after a 2 min dark adapt. For flow cytometry experiments, adherent cells were detached and then resuspended in 1 mL of Dulbecco’s phosphate-buffered saline. GFP fluorescence was measured with a Beckman Coulter Gallios Flow Cytometer using a 488 nm excitation with 525/40 band pass to identify GFP-positive cells. Percent transfection was calculated by dividing the total number of GFP-positive cell counts by the total number of cells counted.

### Single-channel patch clamp recordings

The coding sequences for fetal and adult nicotinic acetylcholine receptor subunits in pRBG4 were provided by Steven Sine (Mayo Clinic). For comparison, each subunit was cloned by restriction digestion (EcoRI) into the pUdOm2.1 backbone for single-channel measurements. AChR cDNAs in pRBG4 or pUdOm1.2 were transfected into BOSC 23 cells [36], originally from ATCC (CRL11270), but provided by Steven M. Sine (Mayo Clinic) (RRID:<u>CVCL_4401</u>). Cells were maintained in Dulbecco’s modified Eagle’s medium (DMEM; Corning) containing 10% (vol/vol) fetal bovine serum (Gibco) at 37°C, until they reached 50–70% confluence. Cells were then transfected using calcium phosphate precipitation, and transfections terminated after 3–4 h by exchanging the medium. Recordings were obtained one day post transfection (16 - 24 hours after exchanging the medium). A separate plasmid encoding GFP was included to facilitate identification of transfected cells. Single-channel patch clamp recordings were performed as previously described [37]. Recordings from BOSC23 cells transiently transfected with muscle-type acetylcholine receptor subunits (⍺, β, δ and γ/ε) in either pRBG4 or pUdOm2.1 plasmid backbones. Recordings were obtained in a cell-attached patch configuration. All recordings were obtained with a membrane potential of –120 mV, with room temperature maintained between 20 and 22°C. The external bath solution contained 142 mM KCl, 5.4 mM NaCl, 0.2 mM CaCl2, and 10 mM 4-(2-hydroxyethyl)–1-piperazineethanesulfonic acid (HEPES), adjusted to pH 7.40 with KOH. The pipette solution contained 80 mM KF, 20 mM KCl, 40 mM K-aspartate, 2 mM MgCl2, 1 mM ethylene glycol-bis(β-aminoethyl ether)-*N*,*N*,*N′*,*N′*-tetraacetic acid, and 10 mM HEPES, adjusted to a pH of 7.40 with KOH. Acetylcholine was added to pipette solutions to a concentration of 30 µM and stored at –80°C. Patch pipettes were fabricated from type 7052 or 8250 non-filamented glass (King Precision Glass) with inner and outer diameters of 1.15 and 1.65 mm, respectively, and coated with SYLGARD 184 (Dow Corning). Prior to recording, electrodes were heat polished to yield a resistance of 5–8 MΩ. Single-channel currents were recorded on an Axopatch 200B patch clamp amplifier (Molecular Devices), with a gain of 100 mV/pA and an internal Bessel filter at 100 kHz. Data were sampled at 1.0 μs intervals using a BNC-2090 A/D converter with a National Instruments PCI 6111e acquisition card and recorded by the program Acquire (Bruxton).

### Mammalian cell line authentication

Approximately 5 million confluent cells were harvested and their total DNA isolated (E.Z.N.A. Tissue DNA Kit), and then submitted to The Centre for Applied Genomics Genetic Analysis Facility (The Hospital for Sick Children, Toronto, Canada) for STR profiling using Promega’s GenePrint 24 System. A similarity search on the 8,389 human cell lines with STR profiles in Cellosaurus release 45.0 was conducted on the resulting STR profiles. The BOSC23 cell line shares closest identity (87.5%, CLASTR 1.4.4 STR Similarity Search Tool score) with cell line Anjou 65 (CVCL_3645). Anjou 65 is a child of CVCL_1926 (HEK293T/17) and is itself a parent line of CVCL_X852 (Bartlett 96). Bartlett 96 is the parent line of BOSC 23 [38, 39]. The HEK293 cell line STR profile, shared 100% identity with HEK293 (CVLC_0045). PCR tests confirmed that the cells were free from detectable mycoplasma contamination [38, 39].

## Author Information

### Corresponding authors

**Adam J. Shuhendler** - Center for Chemical and Synthetic Biology, Department of Chemistry and Biomolecular Sciences, University of Ottawa, Ottawa, ON K1N 6N5, Canada, https://orcid.org/0000-0001-6952-5217

**Corrie J.B. daCosta** - Center for Chemical and Synthetic Biology, Department of Chemistry and Biomolecular Sciences, University of Ottawa, Ottawa, ON K1N 6N5, Canada, https://orcid.org/0000-0002-9546-5331

**François-Xavier Campbell-Valois** - Center for Chemical and Synthetic Biology, Department of Chemistry and Biomolecular Sciences, University of Ottawa, Ottawa, ON K1N 6N5, Canada, https://orcid.org/0000-0001-5105-2968;

### Authors

**France Ourida Manigat** - Center for Chemical and Synthetic Biology, Department of Chemistry and Biomolecular Sciences, University of Ottawa, Ottawa, ON K1N 6N5, Canada, https://orcid.org/0009-0007-2365-0021

**Louise B. Connell** - Center for Chemical and Synthetic Biology, Department of Chemistry and Biomolecular Sciences, University of Ottawa, Ottawa, ON K1N 6N5, Canada, https://orcid.org/0000-0002-3276-8626

**Brittany N. Stewart** - Center for Chemical and Synthetic Biology, Department of Chemistry and Biomolecular Sciences, University of Ottawa, Ottawa, ON K1N 6N5, Canada, https://orcid.org/0009-0002-2565-3748

**Abdel-Rahman LePabic** - Center for Chemical and Synthetic Biology, Department of Chemistry and Biomolecular Sciences, University of Ottawa, Ottawa, ON K1N 6N5, Canada, https://orcid.org/0009-0004-4937-5919

**Christian J.G. Tessier** - Center for Chemical and Synthetic Biology, Department of Chemistry and Biomolecular Sciences, University of Ottawa, Ottawa, ON K1N 6N5, Canada, https://orcid.org/0000-0002-6006-7755

**Johnathon R. Emlaw** - Center for Chemical and Synthetic Biology, Department of Chemistry and Biomolecular Sciences, University of Ottawa, Ottawa, ON K1N 6N5, Canada, https://orcid.org/0000-0001-7255-182X

**Nicholas D. Calvert** - Center for Chemical and Synthetic Biology, Department of Chemistry and Biomolecular Sciences, University of Ottawa, Ottawa, ON K1N 6N5, Canada, https://orcid.org/0000-0002-6637-9063

**Anthony Rössl** - Center for Chemical and Synthetic Biology, Department of Chemistry and Biomolecular Sciences, University of Ottawa, Ottawa, ON K1N 6N5, Canada, https://orcid.org/0000-0002-2496-6358

### Author Contribution

F.X.C.V. designed the backbone of pUdO#a,c,ks,te,z, F.O.M. constructed and characterized this plasmid set and performed all experiments with *Shigella*. F.X.C.V., F.O.M. and A.R. designed and constructed the pTSAR3.#t. B.S. and A.R.L. constructed the pUdOm#.# series. L.B.C. characterized the pUdOm#.#, and N.D.C. assisted with flow cytometry. J.R.E. and C.J.G.T. performed patch clamp experiments. F.O.M., L.B.C., C.J.B.d.C. and F.X.C.V. wrote the manuscript. F.X.C.V., C.J.B.d.C. and A.J.S. conceived and supervised the project.

### Notes

The authors declare no competing financial interest.

## Acknowledgements

F.X.C.V. acknowledges funding of this project by the grant No. 05587 from the Natural Sciences and Engineering Research Council (NSERC), the grant No. 34789 from the Canada Foundation for Innovation (CFI), and the grant No. 159517 from the Canadian Institutes of Health Research. C.J.B.d.C. acknowledges grants from the NSERC (RGPIN-2016-04801), the CFI (34475), the CIHR (377068), as well as a New Frontiers in Research Fund-Exploration Grant (NFRFE-2018- 00064). A.J.S. acknowledges support from the Government of Ontario Early Researcher Award. The research office and the Faculty of Science of the University of Ottawa contributed funds to the bioGARAGE. F.O.M. is a doctoral postgraduate NSERC scholar. We are grateful to Jeffrey W. Keillor, John P. Pezacki, and Christopher N. Boddy for access to equipment.

