## Supplementary material for "pUdOs: concise plasmids for bacterial and mammalian cells": Figures S1-S3 and Tables S1-S3

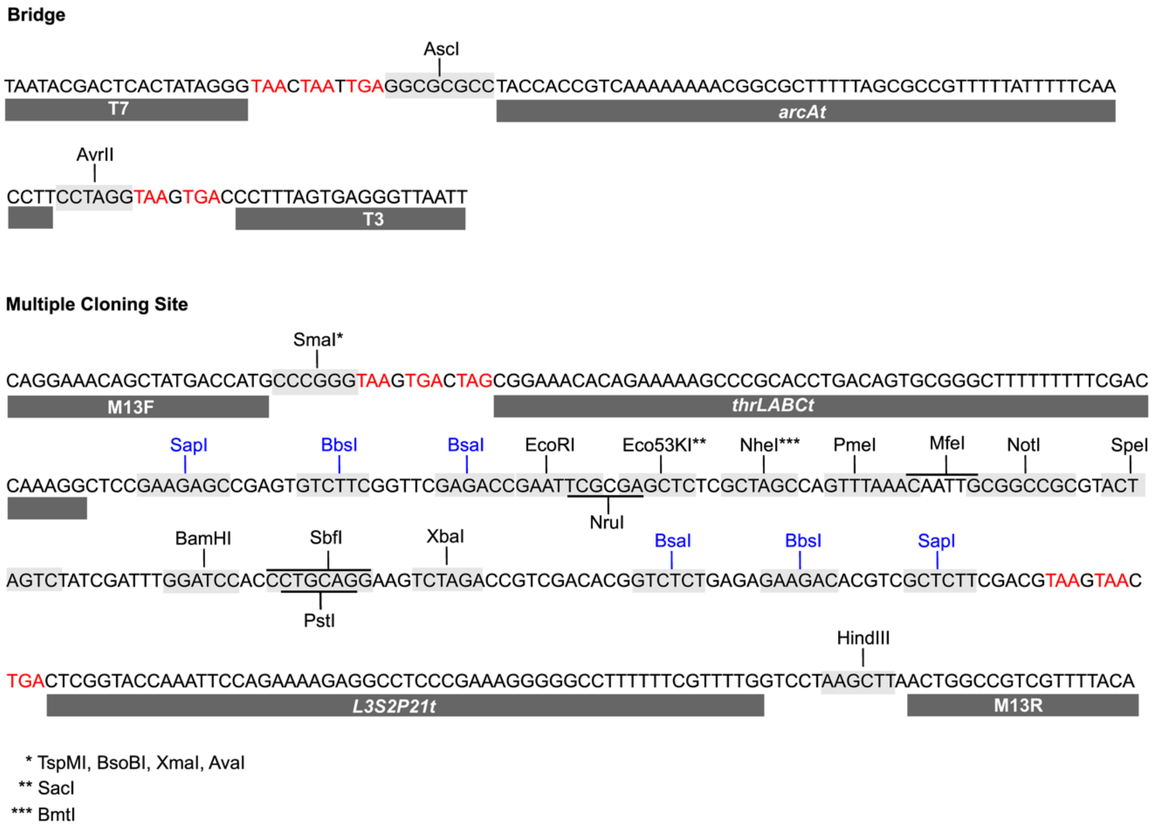


**Figure S1. Sequence of MCS and Bridge fragments.** Restriction sites for subcloning are shown. Type IIS restriction sites for Golden Gate are highlighted in blue and STOP codons are in red.

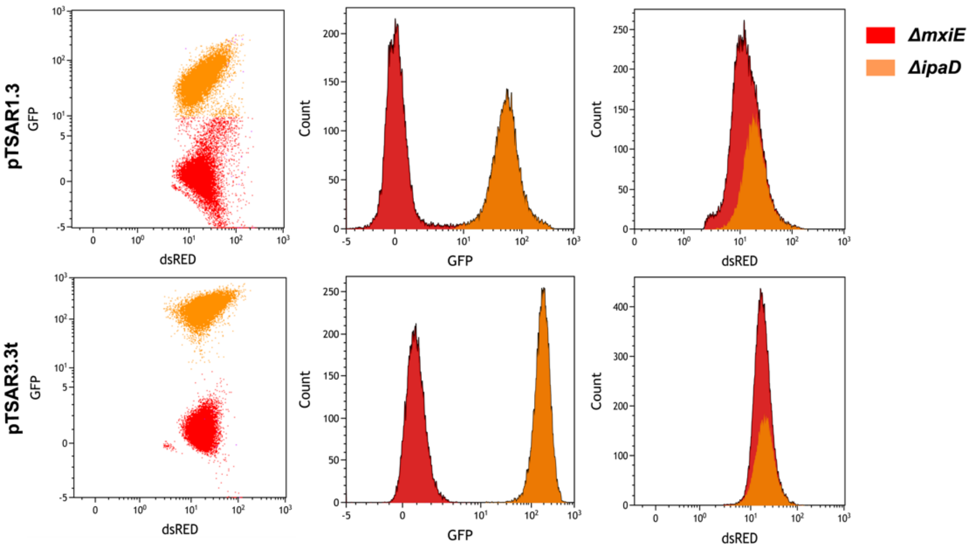


**Figure S2. FC dot plots comparing pTSAR1.3 to pTSAR3.3t GFP and RFP fluorescence in *ΔipaD* and *ΔmxiE* bacteria**.

| pUdOm1.1 | pUdOm1.2 | pUdOm1.3 | pUdOm1.4 | pUdOm1.5 | pUdOm2.1 | pUdOm2.2 | pUdOm2.3 | pUdOm2.4 | pUdOm2.5 | pUdOm3.1 | pUdOm4.1 | pUdOm5.1 | pUdOm6.1 | pUdOm7.1 | pUdOm8.1 | pcDNA3.1 | pcDNA3.0 | pRBG4 | |
|---|---|---|---|---|---|---|---|---|---|---|---|---|---|---|---|---|---|---|---|
| | >0.9999 | **0.0005** | **<0.0001** | **<0.0001** | >0.9999 | 0.8170 | **<0.0001** | **<0.0001** | **<0.0001** | >0.9999 | 0.9659 | >0.9999 | >0.9999 | 0.6290 | 0.9580 | >0.9999 | >0.9999 | >0.9999 | pUdOm1.1 |
| | | **0.0007** | **<0.0001** | **<0.0001** | >0.9999 | 0.8519 | **<0.0001** | **<0.0001** | **<0.0001** | >0.9999 | 0.9519 | >0.9999 | >0.9999 | 0.6761 | 0.9706 | >0.9999 | >0.9999 | >0.9999 | pUdOm1.2 |
| | | | **0.0075** | 0.0625 | **0.0157** | 0.2438 | >0.9999 | **0.0001** | **0.0012** | **0.0113** | **<0.0001** | **0.0013** | **<0.0001** | 0.4040 | 0.1104 | **0.0005** | **0.0011** | **0.0002** | pUdOm1.3 |
| | | | | >0.9999 | **<0.0001** | **<0.0001** | **0.0476** | 0.9994 | >0.9999 | **<0.0001** | **<0.0001** | **<0.0001** | **<0.0001** | **<0.0001** | **<0.0001** | **<0.0001** | **<0.0001** | **<0.0001** | pUdOm1.4 |
| | | | | | **<0.0001** | **<0.0001** | 0.2836 | 0.9028 | 0.9985 | **<0.0001** | **<0.0001** | **<0.0001** | **<0.0001** | **<0.0001** | **<0.0001** | **<0.0001** | **<0.0001** | **<0.0001** | pUdOm1.5 |
| | | | | | | 0.9998 | **0.0003** | **<0.0001** | **<0.0001** | >0.9999 | 0.3964 | >0.9999 | 0.9439 | 0.9964 | >0.9999 | >0.9999 | >0.9999 | 0.9986 | pUdOm2.1 |
| | | | | | | | **0.0154** | **<0.0001** | **<0.0001** | 0.9993 | **0.0290** | 0.9270 | 0.2937 | >0.9999 | >0.9999 | 0.8079 | 0.9089 | 0.6539 | pUdOm2.2 |
| | | | | | | | | **0.0009** | **0.0088** | **0.0002** | **<0.0001** | **<0.0001** | **<0.0001** | **0.0371** | **0.0046** | **<0.0001** | **<0.0001** | **<0.0001** | pUdOm2.3 |
| | | | | | | | | | >0.9999 | **<0.0001** | **<0.0001** | **<0.0001** | **<0.0001** | **<0.0001** | **<0.0001** | **<0.0001** | **<0.0001** | **<0.0001** | pUdOm2.4 |
| | | | | | | | | | | **<0.0001** | **<0.0001** | **<0.0001** | **<0.0001** | **<0.0001** | **<0.0001** | **<0.0001** | **<0.0001** | **<0.0001** | pUdOm2.5 |
| | | | | | | | | | | | 0.4722 | >0.9999 | 0.9681 | 0.9913 | >0.9999 | >0.9999 | >0.9999 | 0.9996 | pUdOm3.1 |
| | | | | | | | | | | | | 0.8926 | >0.9999 | **0.0118** | 0.0818 | 0.9688 | 0.9128 | 0.9934 | pUdOm4.1 |
| | | | | | | | | | | | | | >0.9999 | 0.7956 | 0.9910 | >0.9999 | >0.9999 | >0.9999 | pUdOm5.1 |
| | | | | | | | | | | | | | | 0.1593 | 0.5372 | >0.9999 | >0.9999 | >0.9999 | pUdOm6.1 |
| | | | | | | | | | | | | | | | >0.9999 | 0.6174 | 0.7637 | 0.4469 | pUdOm7.1 |
| | | | | | | | | | | | | | | | | 0.9544 | 0.9870 | 0.8745 | pUdOm8.1 |
| | | | | | | | | | | | | | | | | | >0.9999 | >0.9999 | pcDNA3.1 |
| | | | | | | | | | | | | | | | | | | >0.9999 | pcDNA3.0 |
| | | | | | | | | | | | | | | | | | | | pRBG4 |

**Figure S3. Significance values for pUdOm luciferase assays**. HEK293 cells were transfected with pUdOm#.# and control plasmids containing the firefly luciferase coding sequence. All calculations were performed on GraphPad Prism using a one-way ANOVA and Tukey's multiple comparison test. Significant p-values ($p<0.05$) are bolded.

### Supplementary Table 1. List of primers

| Primer name | Sequence (5’ – 3’) |
|---|---|
| colE1_F | agtgaccctttagtgagggttaattTAGAAAAGATCAAAGGATCTTCTTGAGATCCTTTTTTTCT[illegible] |
| colE1_R | cgggcatggtcatagctgtttcctgGGCCGCGTTGCTGGCGTT |
| SC101_F | ccctttagtgagggttaattGAGTTATACACAGGGCTG |
| SC101_R | atggtcatagctgtttcctgTCAGATCCTTCCGTATTTAG |
| 15A_F | ccctttagtgagggttaattCTAGAAATATTTTATCTGATTAATAAGATG |
| 15A_R | atggtcatagctgtttcctgCTAGCGGAGTGTATACTG |
| BBR1_F | ccctttagtgagggttaattGCGGCCACCGGCTGGCTC |
| BBR1_R | atggtcatagctgtttcctgCTACCGGCGCGGCAGCGTG |
| AmpR_GA_F | aagcttaactggccgtcgttttacaTTCAAATATGTATCCGCTC |
| AmpR_GA_R | agttaccctatagtgagtcgtattaTTACCAATGCTTAATCAGTG |
| TmpR_GA_F | aagcttaactggccgtcgttttacaCTGTTGACAATTAATCATCGG |
| TmpR_GA_R | agttaccctatagtgagtcgtattaTTAGGCCACACGTTCAAG |
| CamR_GA_F | taactggccgtcgttttacaACGTAAGAGGTTCCAACTTTCACCATAATGAAATAAGATCAC[illegible] |
| CamR_GA_R | ccctatagtgagtcgtattaTTACGCCCCGCCCTGCCA |
| ZeoR_GA_F | taactggccgtcgttttacaGTGTTGACAATTAATCATCGGCATAGTATATCG |
| ZeoR_GA_R | ccctatagtgagtcgtattaTCAGTCCTGCTCCTCGGC |
| SmR_GA_F | taactggccgtcgttttacaCCAAGGTTGCCGGGTGAC |
| SmR_GA_R | ccctatagtgagtcgtattaTTATTTGCCGACTACCTTGGTGATC |
| KanR_GA_F | taactggccgtcgttttacaTAGCTTGCAGTGGG |
| KanR_GA_R | ccctatagtgagtcgtattaTCAGAAGAACTCGTCAAG |
| TetR_GA_F | taactggccgtcgttttacaTACGGCCCCAAGGTCCAAAC |
| TetR_GA_R | ccctatagtgagtcgtattaCTAAGCACTTGTCTCCTGTTTACTCC |
| MCS1_GA_S | CAGGAAACAGCTATGACCAT |
| MCS1_GA_R | TGTAAAACGACGGCCAGTTA |
| Bridge_GA_S | TAATACGACTCACTATAGGGTA |
| Bridge_GA_R | AATTAACCCTCACTAAAGGGT |
| Pudo1a_fwd | ACTGACTCGGTACCAAATTCC |
| Pudo1a_rev | CGGAGCCTTTGGTCGAAAAAAAAAG |
| GFPsfm2_fwd | gtttctggtctcccgctagcGAATTGCCCTCGTATTAAATGTGTATCG |
| GFPsfm2_rev | gaatttggtaccgagtcagttctagaTTATGCGGCCAGCGCATAATTTTC |
| GFPsfm2_rev2_[illegible] | gaatttggtaccgagtcagtgttaacTTATGCGGCCAGCGCATAATTTTC |
| L3S3P21_fwd | ccatggagatatcCCAATTATTGAAGGCCTCCCTAACGGGGGGCCTTTTTTTGTTTCTGGT[illegible] |
| L3S3P21_rev | gctagcGGGAGACCAGAAACAAAAAAAGGCCCCCCGTTAGGGAGGCCTTCAATAAT[illegible] |
| rpsM_fwd | tttttcgaccaaaggctccggcggccgcGATTTTTTCGCATATTTTTCTTGC |
| MS165_rev | TAATTGGGATATCTCCATGGCTACAGGAACAGGTGGTG |

| | |
|---|---|
| Cerulean_rev | taattgggatatctccatggttaCTTGTACAGCTCGTCCATG |
| mCherry_rev | taattgggatatctccatggCTACTTGTACAGCTCGTC |

**Supplementary Table 2. List of origin of replication and their characteristics.** Incompatibility groups and copy number of each origin shown are obtained from the literature.

| Origins | Incompatibility group | Copy number | Source plasmid |
|---|---|---|---|
| colE1*(pMB1) | A | 500-700 | pUC18 |
| 15A | B | 10 -12 | pSU2718 |
| SC101 | C | 5 | pKS011 |
| BBR1 | N/A* | ~ 5 | pBT-L1 |

*non- classical

**Supplementary Table 3. List of Selection markers and their source**

| Source plasmid | Parts obtained | Gene name |
|---|---|---|
| pTKDP-dhfr | TmpR | Dihydrofolate reductase |
| pUC18 | AmpR | ß-lactamase |
| pSU2718 | CmR | cml Acetyltransferase |
| pZeo-SV2 | ZeoR | sh ble |
| pRep4 | KanR | APH(3')-II family aminoglycoside O-phosphotransferase (Tn5) |
| pTKLP-tetA | TetR | tetracycline resistance protein, class C (tetA) |
| pTKRED | SmR | aminoglycoside resistance protein (aadA) |
